# Neo-epitope identification by weakly-supervised peptide-TCR binding prediction

**DOI:** 10.1101/2023.08.02.550128

**Authors:** Yuli Gao, Yicheng Gao, Wannian Li, Siqi Wu, Feiyang Xing, Chi Zhou, Shaliu Fu, Guohui Chuai, Qinchang Chen, He Zhang, Qi Liu

**Affiliations:** Key Laboratory of Spine and Spinal Cord Injury Repair and Regeneration (Tongji University), Ministry of Education, Orthopaedic Department of Tongji Hospital, Frontier Science Center for Stem Cell Research, Bioinformatics Department, School of Life Sciences and Technology, Tongji University, Shanghai, China; Translational Medical Center for Stem Cell Therapy and Institution for Regenerative Medicine, Shanghai East Hospital, Frontier Science Center for Stem Cell Research, Bioinformatics Department, School of Life Sciences and Technology, Tongji University, Shanghai, 200092, China; Research Institute of Intelligent Computing, Zhejiang Lab, Hangzhou 311121, China; Shanghai Research Institute for Intelligent Autonomous Systems 201804, China

## Abstract

The identification of T cell neo-epitopes is fundamental and computational challenging in tumor immunotherapy study. As the binding of pMHC - T cell receptor (TCR) is the essential condition for neo-epitopes to trigger the cytotoxic T cell reactivity, several computational studies have been proposed to predict neo-epitopes from the perspective of pMHC-TCR binding recognition. However, they often failed with the inaccurate binding prediction for a single pMHC -TCR pair due to the highly diverse TCR space. In this study, we proposed a novel weakly-supervised learning framework, *i*.*e*., *TCRBagger*, to facilitate the personalized neo-epitope identification with weakly-supervised peptide-TCR binding prediction by bagging a sample-specific TCR profile. *TCRBagger* integrates three carefully designed learning strategies, *i*.*e*. a self-supervised learning strategy, a denoising learning strategy and a Multi-Instance Learning (MIL) strategy in the modeling of peptide-TCR binding. Our comprehensive tests revealed that *TCRBagger* exhibited great advances over existing tools by modeling interactions between peptide and TCR profiles. We further applied *TCRBagger* in different clinical settings, including (1) facilitating the peptide-TCR binding prediction under MIL using single-cell TCR-seq data. (2) improving the patient-specific neoantigen prioritization compared to the existing neoantigen identification tools. Collectively, *TCRBagger* provides novel perspectives and contributions for identifying neo-epitopes as well as discovering potential pMHC-TCR interactions in personalized tumor immunotherapy.

## Background

Tumor immunotherapy by exploring and stimulating the immune system has achieved great success^1^. Personalized vaccines designed to trigger de novo T cell responses against neo-epitopes have demonstrated robust tumor-specific immunogenicity and preliminary evidence of antitumor activity in melanoma and other cancers^2^. HLA-I presented neo-epitopes or neoantigens are peptides presented on the tumor cell surface by human leukocyte antigen (HLA) class I protein, recognized by CD8 T cells^3^ with TCR, eventually eliciting immune response to tumor. With the development of next-generation sequencing (NGS) data at the personalized level, several computational studies for personalized single-nucleotide variant (SNV)-based neo-epitope identification have been developed^4–6^. Generally, these methods use different scoring functions to prioritize tumor specific peptides by considering different features, such as peptide expression level, mutation allele fraction or binding affinity of HLA-peptide pairs^7^. However, few candidate neo-epitopes identified by the existed algorithms were able to elicit the CD8 T cell response in vitro or in vivo ^8–10^ as they failed to accurately model the recognition between neopeptides and tumor-infiltrating T cells. The identification of neo-epitopes that can elicit T cell response is fundamental and computational challenging in immunology study^11^.

Although several studies have been proposed to predict tumor neo-epitopes from the perspective of pMHC-TCR binding recognition^12–14^, they often failed due to the inaccurate binding prediction for a single pMHC -TCR pair due to the highly diverse TCR space. To this end, several machine learning models, including Gaussian process^15^ and deep neural network models have been presented for single TCR-peptide binding prediction^13,16–18^, while these methods achieved limited performance on unseen peptide-TCR pairs in general. Accurately identifying the tumor specific neo-epitope by studying the interaction between peptide and patient-specific tumor-infiltrating T cells are still lack.

In this study, we aim at identifying the HLA-I presented neo-epitopes that could elicit CD8 T-cell immune response based on the simplified assumptions that (1) the binding of pMHC - TCR is the essential condition for HLA-I presented neo-epitope to trigger the CD8 T-cell response, and (2) the HLA-I presented neo-epitopes are recognized or bound by at least one TCR in the T cell response, while those peptides failed to trigger the CD8 T-cell response are not recognized by any TCR. Therefore, there does not necessarily require to model the interaction between the neo-epitope and a single TCR explicitly in the identification of neo-epitope, rather an interaction recognition between the neo-epitope and the sample-specific TCR profile is needed. To this end, we present a novel weakly-supervised learning framework, *i*.*e*., *TCRBagger*, which aims to identify neo-epitope by considering its interaction with a sample-specific TCR profile rather than a single TCR in the tumor specific T cell response, considering that the exactly binding TCRs for the neo-epitopes are often unknown and difficult to predict in clinic. *TCRBagger* integrates three carefully designed learning strategies in peptide-TCR interaction modeling, including (1) a self-supervised learning strategy to learn the context-based representation of peptides presented by major histocompatibility complex (MHC) class I molecules (2) a denoising learning strategy to weaken the impact of incorrectly experimental annotated immunogenic peptides in the modeling, and finally (3) a multi-instance learning (MIL) strategy to predict the interaction between peptide and the bagging of a sample-specific TCR profile rather than a single TCR, which avoids the inaccuracy in the recognition of a single peptide-TCR interaction. Our comprehensive tests revealed that *TCRBagger* exhibited great advances over existing tools by modeling interactions between peptide and TCR profiles. We further applied *TCRBagger* in different clinical settings, including (1) facilitating the peptide-TCR binding prediction under MIL using single-cell TCR-seq data. (2) improving the neoantigen prioritization compared to the existing neoantigen identification tools including MuPeXI^6^, Neopepsee^5^ and pTuneos^4^.

## Results

### 1. The general framework of *TCRBagger*

*TCRBagger* was developed with the goal of identifying neo-epitopes with peptide-TCR binding prediction by bagging a sample-specific TCR profile instead of a single TCR in the T cell response (**see Methods**). The basic idea of *TCRBagger* is inspired by a kind of weakly supervised learning, *i*.*e*., a multi-instance learning (MIL) model^19^, which was originally proposed to solve the problem of drug activity prediction due to the coexistence of “conformations” in each molecule^20^. In MIL, training samples are arranged in sets, called bags, and a label is provided for the entire bag, and the individual labels of the sample contained in the bags are not provided. In our study, MIL can alleviate the labeling burden for a single peptide-TCR binding, since they are often unknown and difficult to obtain or predict in clinic. In this case, the individual binding labels of TCRs to a given peptide are not explicitly provided. We only label the peptide as immunogenic or not by arranging the sample-specific TCR profile as a bag. Such weak supervision information is generally obtained more efficiently (**see Methods**). In this study, we clearly define the HLA-I presented immunogenic peptides (neo-epitopes) as those peptides recognized or bound by at least one TCR and trigger the CD8 T cell response, while the non-immunogenic ones are those peptides not recognized by any TCR and failed to trigger the CD8 T-cell response. Of note, several studies have been presented for immune repertoire classification by linking TCR repertories to specific phenotype using MIL^21–23^, however, the explicit study of identifying neo-epitopes by interrogating the interaction of peptide and TCR profile using MIL is still lacking.

In order to learn the intrinsic features of neo-epitopes determined by the sample-specific TCR profile, we systematically integrated various data sources to curate the following three datasets for *TCRBagger* modeling (**see Supplementary Methods**), including (1) ***Peptide immunogenicity dataset***: the HLA-I restricted peptides with experimental annotated immunogenicity label, *i*.*e*., immunogenic or not. These peptides were collected from IEDB, while they lacked the exact TCR binding annotation. Therefore, we used these peptides in the self-supervised pre-training to learn an efficient representation of HLA-I-restricted peptides. We further adopted the denoising learning strategy for these peptides to exclude the potential mislabeled peptides using the learned representation from the self-supervised pre-training. (2) ***Peptide-TCR binding dataset***: the known peptide-TCR binding pairs integrated from four databases, i.e., IEDB^24^, VDJdb^25^, PIRD^26^ and McPas-TCR^27^. Those binding TCRs for the specific peptide were defined as the peptide-specific TCRs, taken as the positive instances sampled for bag construction in MIL in the subsequent analysis. (3) ***Negative control TCR dataset***. These were the TCRs collected from 587 healthy volunteers’ peripheral blood using a multiplexed polymerase chain reaction assay that targets the variable region of the rearranged TCRβ locus^28^, which were taken as the negative instances for bag construction in *TCRBagger* modeling, considering the very low probability of encountering binding TCRs when randomly sampling a part of TCRs from this large pool^18^.(**Fig.1**)

**Fig. 1.**
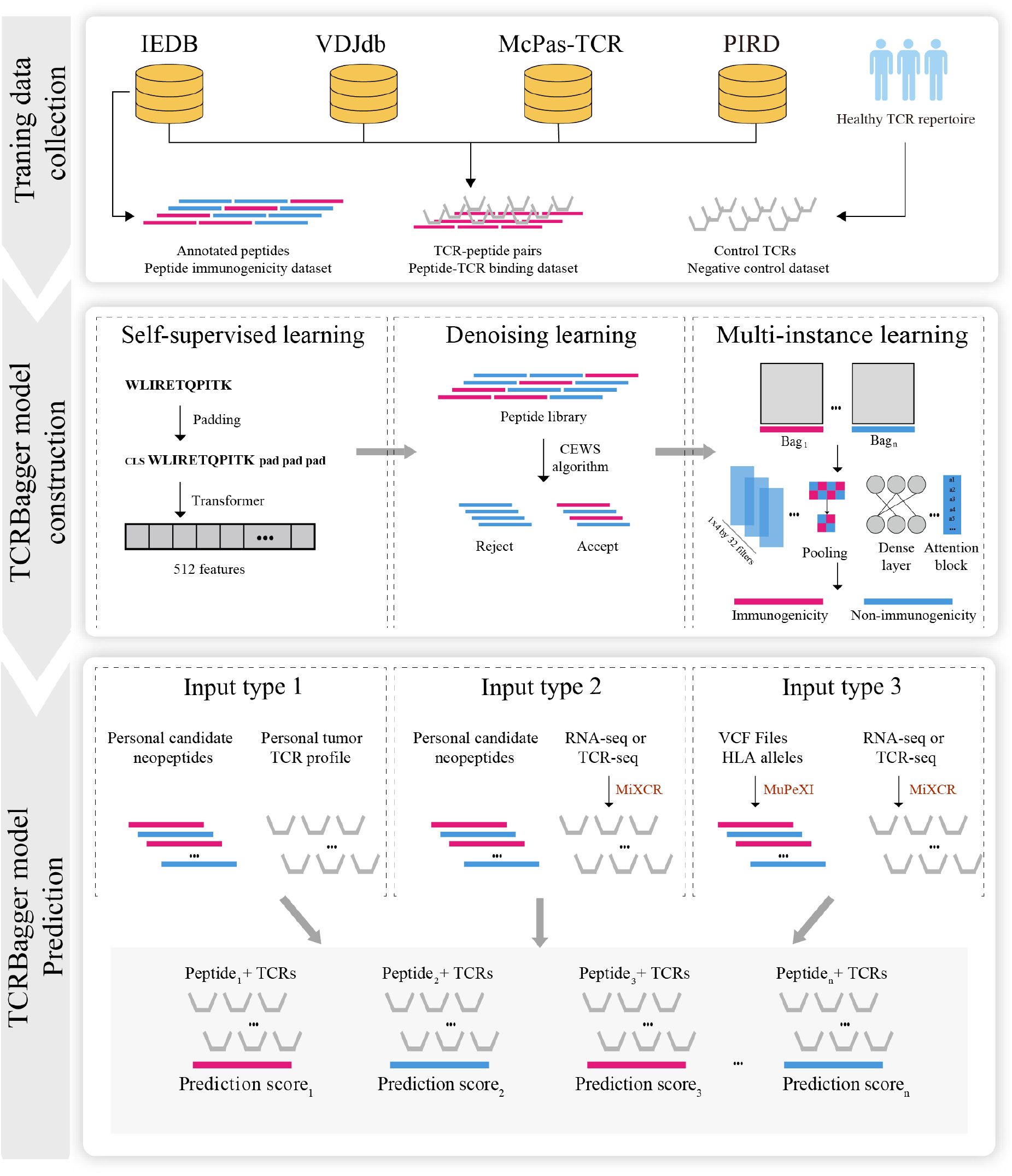
**The general framework of *TCRBagger*. The computational framework of *TCRBagger* is divided into three parts. (1) Training data collection: all datasets for constructing bag-level *TCRBagger* model training are generated from four databases and 587 healthy volunteers’ TCR repertoire. (2) Model construction: three learning algorithms are carefully designed in *TCRBagger*, including *self-supervised learning, denoising learning* and *multi-instance learning*. (3) Model prediction: *TCRBagger* are developed as one-stop software for various input types, including the input of candidate neopeptides and personal tumor TCR profiles, the input of candidate neopeptides and tumor RNA-seq or TCR-seq, and the input of VCF files, HLA alleles and tumor RNA-seq or TCR-seq. *TCRBagger* provides a prediction score for each peptide that represent the probability to be a neo-epitope by modeling peptide recognition with the given TCR profile**.

In order to evaluate the effectiveness of *TCRBagger*, three carefully designed learning strategies were used for identifying neo-epitopes based on the interaction between peptide and the sample-specific TCR profile. First, to learn the context-based representation of HLA-I restricted peptides, we applied a self-supervised learning strategy, which was used in the protein representation area ^29–31^. Second, to exclude label noise in experimentally annotated peptides, we used the cut edge weight statistic (CEWS) algorithm^32^, a denoising learning strategy that considers the situation in which the supervision information is full of noise in peptide annotations ^33^. Of note, the peptide embedding applied in the CEWS analysis is based on the representation obtained by the self-supervised learning in the 1^st^ strategy. Finally, we identify the candidate neo-epitopes by considering the interaction with the sample-specific TCR profile by MIL. (**Fig. 1, See Methods**)

We further developed *TCRBagger* as a user-friendly and one-stop software with various input types provided by users, and output the prediction score for each peptide, representing the probability to be a neo-epitope by considering the sample-specific TCR profile (**Fig.1, See Methods**).

### 2. Comprehensive data exploration demonstrates the rationale of using MIL for peptide-TCR binding modeling

We first comprehensively explored the peptide-TCR interaction pairs in the ***Peptide-TCR binding dataset***, obtaining a total of 666 different peptides with specific TCR bindings with the filter criteria (**see Supplementary Methods, Supplementary Fig.1a, Supplementary Data 1**). Nearly 80% of these peptides were found to bind to 1-10 TCRs, 15% of them were found to bind to 10-100 TCRs and 5% of them were found to bind more than 100 TCRs (**Supplementary Fig.1b**). Then, only 7.3% of immunogenic peptides (positive samples) in the ***Peptide immunogenicity dataset*** own their specific TCR binding information in the ***Peptide-TCR binding dataset***, showing the sparsity of specific TCR binding annotations for most of the peptides (**Supplementary Fig.2c**), which has inspired us to apply MIL to identify neo-epitopes through bagging TCR profiles instead of single TCR-peptide binding.

We also investigated TCR motifs among the TCRs in the ***Peptide-TCR binding dataset*** and the ***Negative control TCR dataset*** using Logomaker ^34^. We found that the lengths of 95% TCRs in those two sets are in the range of 10-20 (**Supplementary Fig.1c**). We further discovered that the leading four sites and the tailing four sites are more conservative than the other sites in both of the two sets (**Fig S1d, e**). These results reveal that the leading and tailing four sites contribute less to the diversity of TCR recognition, reveling these sites may not participate in peptide interaction, which is consistent with previous studies^18,25^.

### 3. Inaccurately experimentally annotated peptides exist in Peptide immunogenicity dataset

We observed 7 IEDB annotated non-immunogenic peptides (negative samples) in the ***Peptide immunogenicity dataset*** have TCR binding information in ***Peptide-TCR binding dataset***, meaning that noise existed among these curated dataset (**Supplementary Fig.2b**).To address this issue, we adopted CEWS algorithm to exclude possible incorrectly experimentally annotated peptides in the ***Peptide immunogenicity dataset***^35^. First, we applied transformer encoder based self-supervised learning to obtain the 512 embedding features for each HLA-I-restricted peptides^29^ (**see Methods, Supplementary Fig.3-4**). Then CEWS algorithm gave each peptide a mislabeled score and took a peptide as mislabeled peptide when that score was higher than 2.5758, which represents the right unilateral *p* − *value* score of normal distribution function with a *p* − *value* of 0.01(**Fig. 2a, see Methods**).

**Fig. 2.**
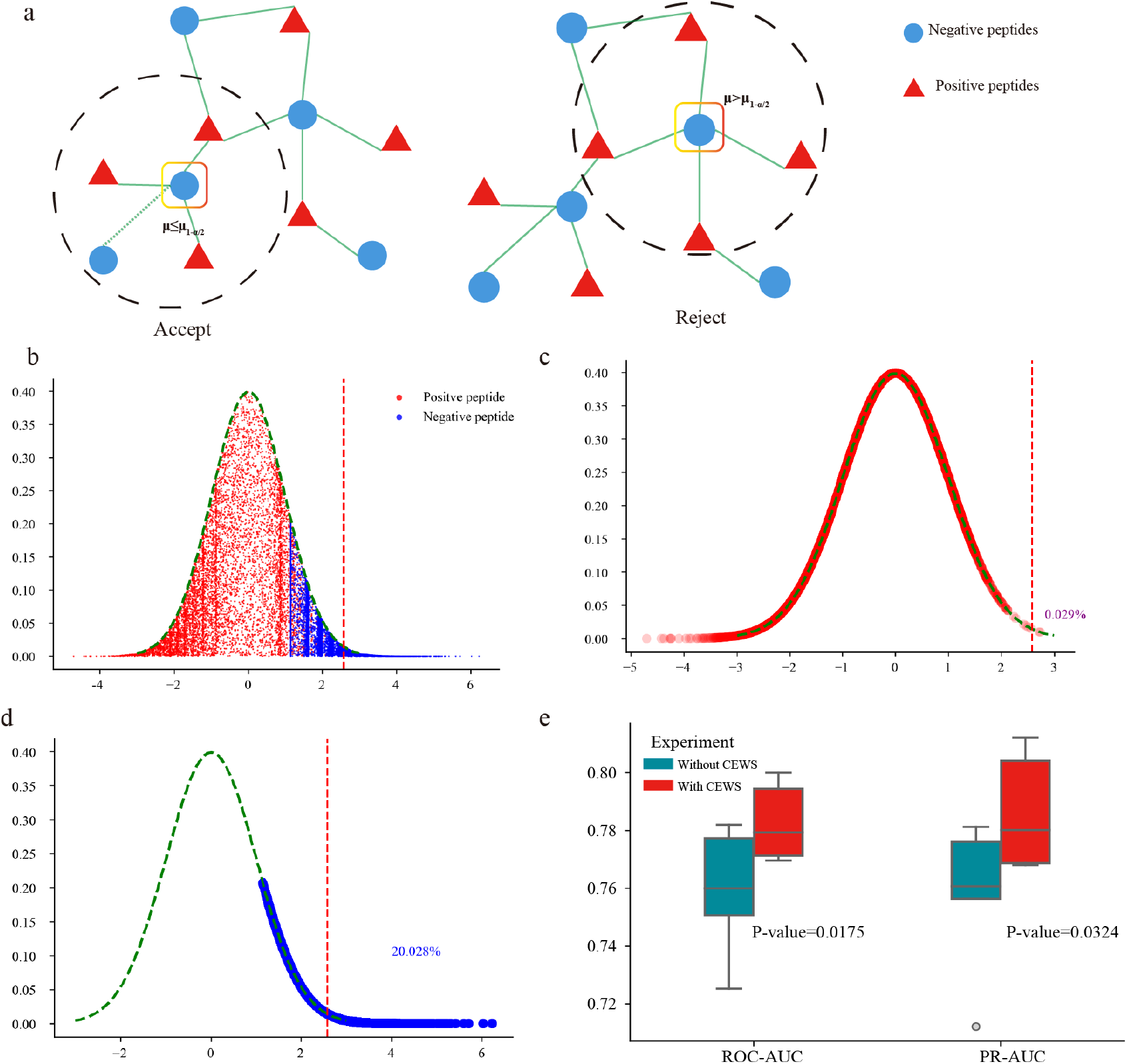
**Illustration of denoising learning strategy. a: The CEWS algorithm identifies the incorrectly annotated peptides. b: The distribution of mislabeled scores given by the CEWS algorithm between positive peptides and negative peptides. The x-axis represents the mislabeled score for each peptide given by CEWS, and the y-axis represents each random height value for each peptide with a normal distribution. c: 99.971% of positive peptides were accepted, and 0.029% were refused by the CEWS algorithm d: 79.972% of negative peptides were accepted, and 20.028% were refused by the CEWS algorithm. e: The improvement of *TCRBagger* after denoising in terms of ROC-AUC and PR-AUC**

As a result, it is shown that the mislabeled scores of positive peptides were smaller than negative peptides, indicating the effectiveness of the CEWS algorithm, as incorrectly annotated negative peptides often occurred with greater possibilities (**Fig. 2b, Supplementary Data 2**). In this case, 0.029% positive peptides with no identified binding TCRs at present were considered as possible incorrectly annotated peptides by CEWS(**Fig.2c**). The lower noise existed in the positive peptide dataset is acceptable, which may come from experimental annotation error. On the other hand, CEWS algorithm also recognized 20.028% of negative peptides as mislabeled peptides (**Fig.2d**). Eventually, we excluded those identified mislabeled peptides by CEWS in the ***Peptide immunogenicity dataset***, which was proven to significantly improve the performance of *TCRBagger* as shown in the subsequent study.

### 4. *TCRBagger* facilitates HLA-I-restricted neo-epitope identification

#### 4.1 The single TCR-peptide binding prediction methods failed on unseen peptides

To illustrate the limitations of traditional single TCR-peptide predictions, we evaluated three state-of-the-art tools, including pMTnet^13^, ERGO-II^14^ and DLpTCR^12^, on the seen peptides and unseen peptides. Seen peptides means those peptides with their binding TCRs are seen in the training of those tools, and we want to predict the binding of new TCRs for these seen peptides. While unseen peptides mean those peptides are totally new in the prediction and they are unseen in the training of those tools. Of note, we used the negative peptides in the ***Peptide immunogenicity dataset*** after CEWS denoising with randomly sampled TCRs from the ***negative control TCR dataset*** to balance the positive TCR-peptide pairs (**Supplementary Data 3**). As a result, pMTnet achieved the best performance in terms of ROC-AUC and PR-AUC, while ERGO-II and DLpTCR all achieved worse performance (**Fig.4a,b)**. Specifically, the performance of pMTnet deceased for unseen peptides (ROC-AUC=0.65, PR-AUC=0.69, **Fig.4a,b**), indicating the limitation of current TCR-peptide prediction tools on the unseen peptide-TCR pairs recognition due to the complicated interaction mechanism of TCR-peptide, while the recognition of the binding specificity for neo-epitope that previously unseen by the immune system is crucial.

#### 4.2 *TCRBagger* facilitates neo-epitope identification with weakly-supervised peptide-TCR binding prediction

The basic idea of *TCRBagger* for neo-epitope identification is to interrogate the interaction between peptide and the sample-specific TCR profile rather than a single TCR with the MIL framework. In this framework, a positive bag means that a peptide in this bag can be bound with at least one TCR, while a negative bag does not. The bag can be taken as a representation of the sample-specific TCR profile. The features of each instance, *i*.*e*., a peptide-TCR pair in a bag, consist of three parts. (1) 75 features encoding a peptide sequence where each amino acid uses five numbers represented by the Atchley factors^36,37^. (2) 100 features encoding a TCR sequence that also uses Atchley factors. (3) 24 features representing the binding properties of the TCR loop with peptides. To this end, amino acid pairwise contact potential (AACP) scales were downloaded from the AA Index database^38^, and the “global-local” algorithm was applied to obtain the optimal binding features of each peptide-TCR instance (**Fig.3a, see Methods**). In addition, we used a gate attention block in *TCRBagger* instead of the traditional MIL pooling layers, such as max pooling, mean pooling and log-sum-exp (LSE) to combine instance-level features into a bag-level feature representation (**Fig.3b, see Methods**). Finally, we constructed 13,100 positive bags using peptides with their randomly sampled binding TCRs in ***Peptide-TCR binding dataset*** and the randomly sampled negative control TCRs in ***Negative control TCR dataset*** to ensure the TCRs number reach to 100 in each bag. In addition, we also build 14,109 negative bags using the peptides in ***peptide-TCR binding dataset*** with their randomly sampled 100 negative control TCRs as well as the negative peptides in ***Peptide immunogenicity dataset*** with their randomly sampled 100 negative control TCRs. Furthermore, two data splitting strategies were applied for testing. In the first splitting scenario, bags were randomly split into five sub-datasets, and the peptides used in the training may be seen in the testing, but their corresponding TCR profiles in each bag were different. It should be noted that this splitting strategy was similar as most TCR-peptide prediction tools applied. In the second splitting scenario, bags were split into five sub-datasets according to peptides in bags, ensuring a more rigorous test that the peptides and TCRs are all unseen to the training dataset (**Fig.4c**).

**Fig. 3.**
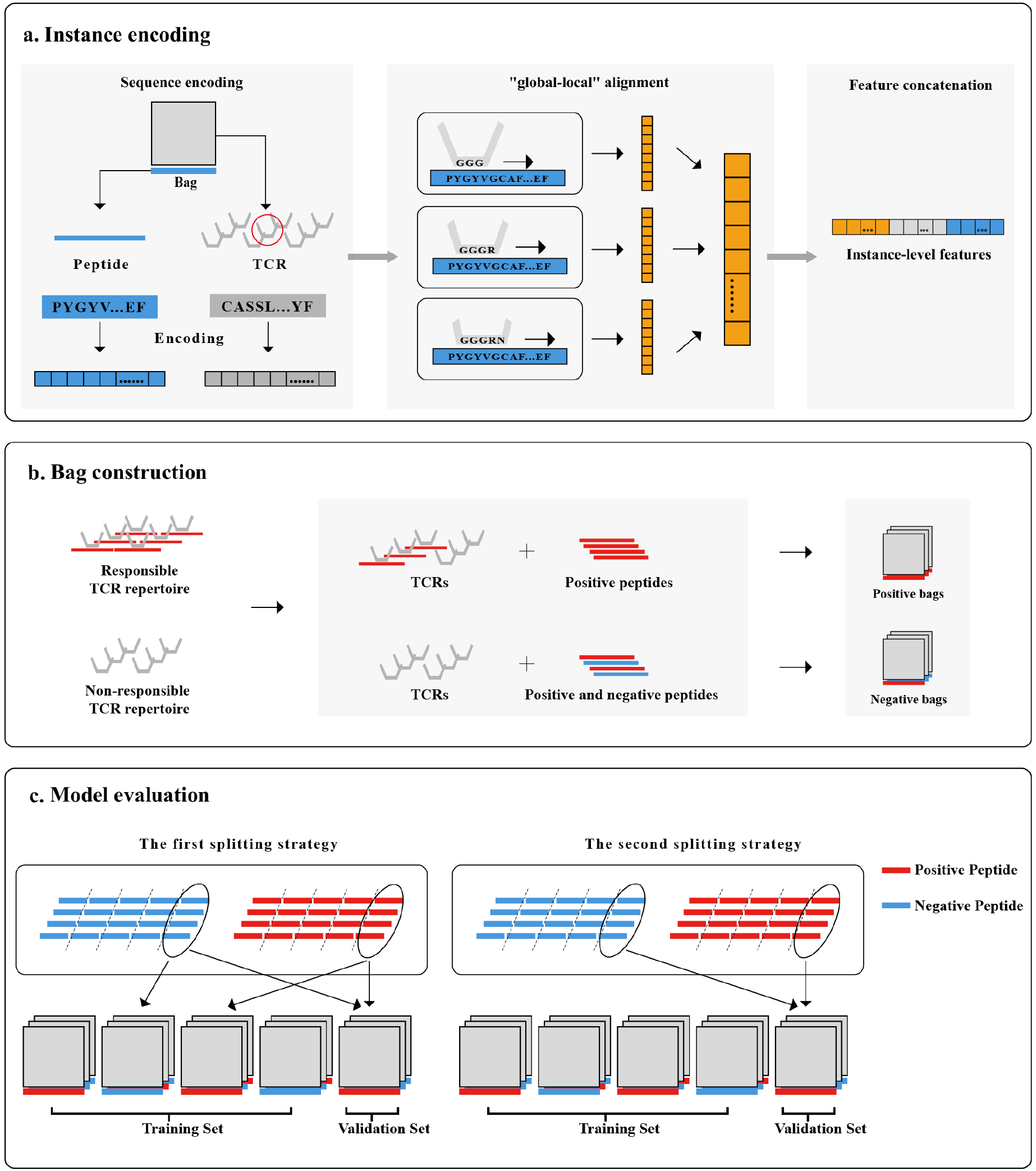
**Overview of bag generation of *TCRBagger*. a. Instance encoding consists of three parts. First, we used the Atchley factor to encode each peptide and TCR sequence. Second, “Global-local” alignment was applied to calculate the binding features of peptides and TCRs. Third, we concatenated the three-part features obtained. b. Simulated bag construction in the test. c. Fivefold model cross-validation of two splitting strategies were performed**.

**Fig. 4.**
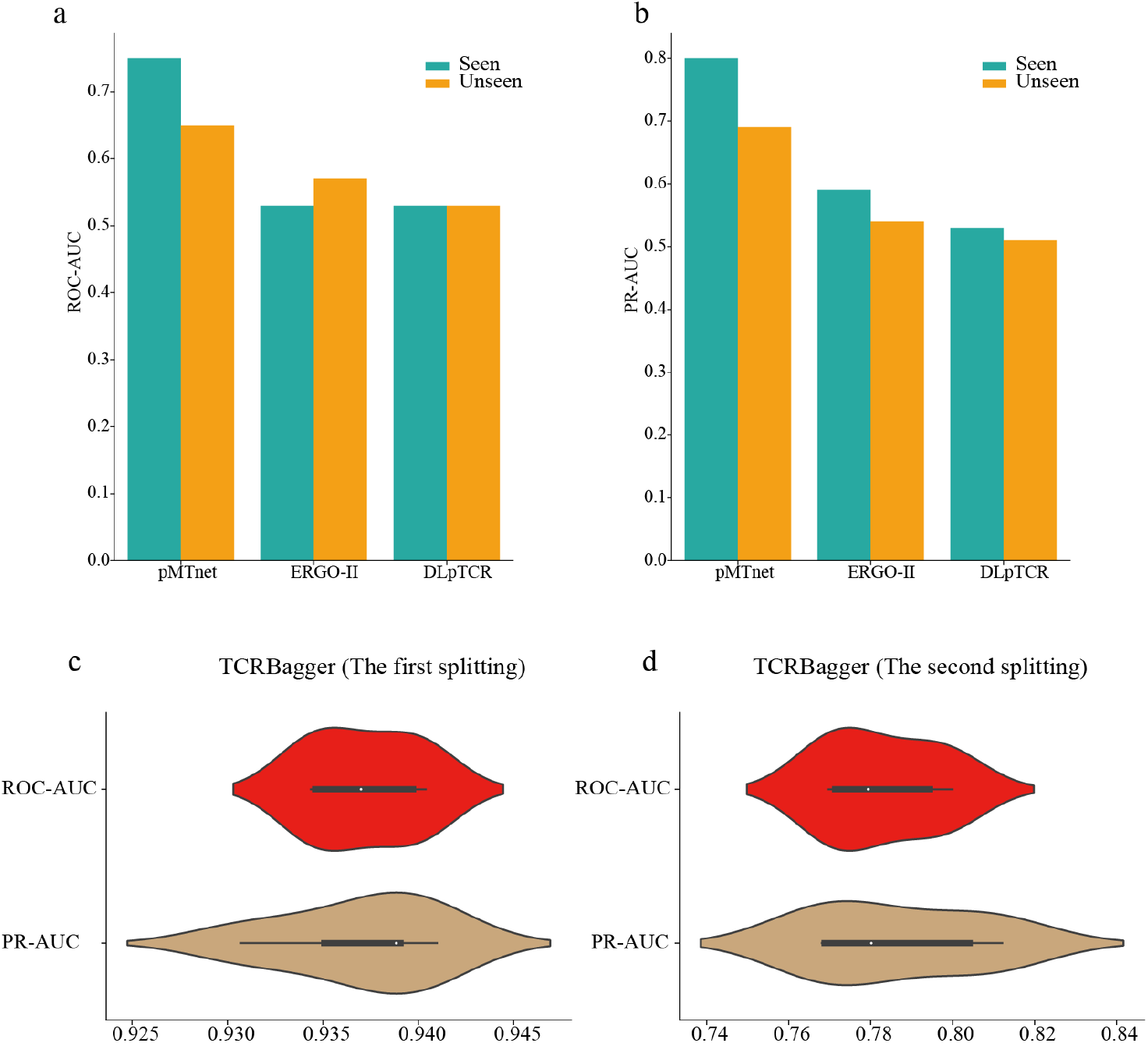
**The performance of *TCRBagger* and the other existing tools. a. The ROC-AUC of pMTnet, ERGO-II, DLpTCR on their seen and unseen dataset. b. The PR-AUC of pMTnet, ERGO-II, DLpTCR on their seen and unseen dataset. c. The ROC-AUC and PR-AUC of *TCRBagger* in the first data splitting scenario with 5-fold cross-validation. d. The ROC-AUC and PR-AUC of *TCRBagger* in the second data splitting scenario with 5-fold cross-validation**.

As a result, the average ROC-AUC and PR-AUC reached 0.938 and 0.937, respectively, in the first data splitting scenario, showing that *TCRBagger* performs quite well in seen peptides by bagging different TCRs (**Fig.4c**). The average ROC-AUC and PR-AUC reached 0.783 and 0.787, respectively, in the second data splitting scenario (**Fig.4d**), meaning that *TCRBagger* also shows good performance on neo-epitope identification for unseen peptides, which is crucial as neo-epitopes are previously unseen for the immune system.

#### 4.3 CEWS algorithm improves the performance of *TCRBagger* by excluding incorrectly annotated peptides

In order to illustrate the importance of denoising in our study, we performed an ablation study to indicate the improvement of the *TCRBagger* performance by excluding incorrectly experimentally annotated peptides in ***Peptide immunogenicity dataset***. To this end, 13,100 positive bags were built using peptides in ***Peptide-TCR binding dataset*** with their randomly sampled binding TCRs and negative control TCRs. 14,005 negative bags were built using negative peptides in ***Peptide immunogenicity dataset*** without excluding the possible incorrectly experimental annotated peptides (*TCRBagger*-without-CEWS). This is compared with the former test using the second splitting strategy, which is denoted as *TCRBagger*-with-CEWS. As a result, *TCRBagger*-with-CEWS performed better, with a 3.15% improvement in ROC-AUC (p value=0.0175) and a 3.87% improvement in PR-AUC (p value=0.0324) compared to *TCRBagger*-without-CEWS, showing the superiority and necessity to exclude label noise in peptide-TCR binding prediction under MIL (**Fig.2e**).

### 5. *TCRBagger* facilitates the neo-epitope identification in single-cell TCR-seq

Single-cell TCR-seq has become increasing popular in immune study, and we evaluate whether *TCRBagger* can be generalized to predict neo-epitopes with TCRs information obtained in single-cell TCR-seq data. Therefore, we collected six datasets from six single-cell TCR-seq experiments using Tetramer-associated T cell receptor sequencing (TetTCR-Seq), linking sample-specific TCR sequences to their congregating antigens in single cells at high throughput ^39^. The CD8 T cells of the six experiments were taken from six human peripheral blood samples. In addition, for each T cell, they used molecular identifier (MID) to obtain the top 5 ranked peptides that are classified as positively binding where “0” indicates that no positively binding peptide was found (**Supplementary Data 4**).We excluded the TCR-peptides binding records with missing CDR3 sequence chains in each experiment and used these data for bag construction. The detailed bag construction information for each experiment is shown in Supplementary Table 1.

Using *TCRBagger* trained on previous testing with the second splitting as a pre-trained model, we fine-tuned *TCRBagger* on every five of six experiments and tested it on the rest of the experiment. As a result, we obtained an average ROC-AUC and PR-AUC of 0.83 and 0.78, respectively, showing that *TCRBagger* can facilitate neo-epitope identification with peptide-TCR binding prediction under MIL using single-cell TCR-seq data (**Fig.5a, b**).

**Fig. 5.**
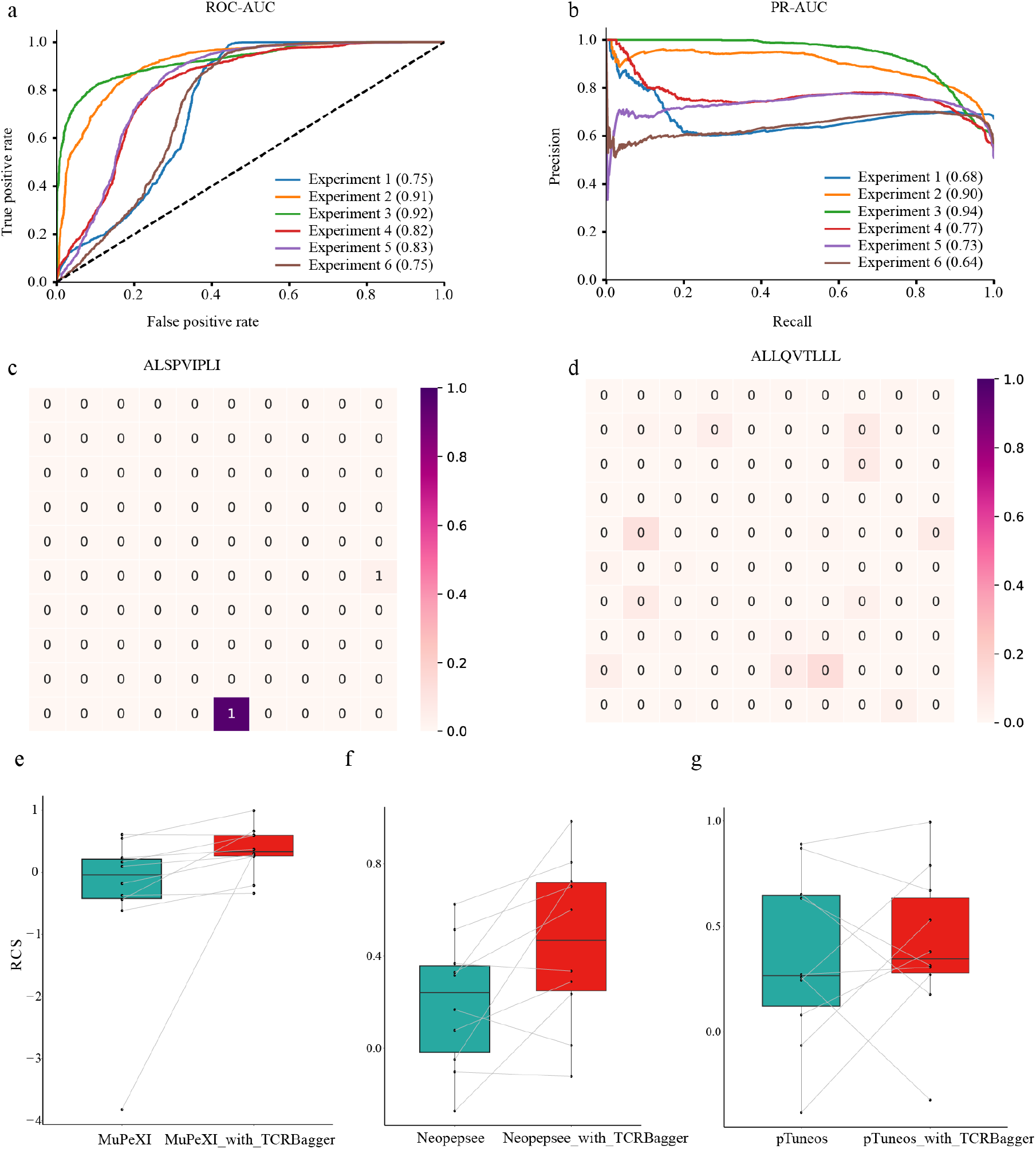
**The performance evaluation of *TCRBaager* on six single-cell experiments and neoantigen identification of ten patients. a: The ROC-AUC of six single-cell experiments. b: The PR-AUC of six single-cell experiments. c: The attention block visualization of a positive bag. A “1” in each square represents a positive TCR-peptide pair. A “0” in each square represents a negative TCR-peptide pair. The darker red represents the greater weights given by the attention block. d: The attention block visualization of a negative bag. A “0” in each square represents a negative TCR-peptide pair. The darker red represents the greater weights given by the attention block. e: *TCRBagger* improves neoantigen identification based on the candidate neoantigen ranking list identified by MuPeXI. f: *TCRBagger* improves neoantigen identification based on the candidate neoantigen ranking list identified by Neopepsee. g: *TCRBagger* improves neoantigen identification based on the candidate neoantigen ranking list identified by pTuneos**.

In this case, the instances, i.e., the peptide-TCR pairs here, contained different amounts of information contributing to bag level prediction, and the attention mechanism instead of the traditional MIL pooling layers was used in *TCRBagger* in the step of combining instance-level features into a bag-level feature representation (**see Supplementary Methods**). Therefore, we could obtain the attention weight for each instance to further understand the key peptide-TCR pair in each bag. Through visualization of the attention, we found that a certain relationship existed between the attention weights and the instance labels from the bag level, and at least one positive instance obtained higher weight in positive bags, indicating that *TCRBagger* is able to capture the fact that there are at least one TCR recognition available for neo-epitope, while negative instances obtained more average weights in negative bags, indicating that none TCR is available to recognize the non-immunogenic ones (**Fig.5c, d, see Methods**). In conclusion, using an attention mechanism, the bag features can be well represented in *TCRBagger*.

### 6. *TCRBagger* substantially improve patient-specific tumor neoantigen prioritization

In order to illustrate that *TCRBagger* can refine the rank of candidate tumor neoantigen identified by the existing tools and enrich the experimentally validated neoantigens on the top, we select three mainstream tumor neoantigen identification tools, including MuPeXI^6^ and Neopepsee^5^ and pTuneos^4^ for comparisons. These three tools do not comprehensively consider the sample-specific T cell response in neoantigen identifications. We applied sequencing data of 10 patients to test their performance, which were also used in previous studies^40–43^. Among them, 25 HLA-I naturally processed and presented neo-peptides were recognized by tumor-infiltrating lymphocytes, meaning these 25 neo-peptides are immunogenic tumor neoantigens. While 22 neo-peptides were not ^4^, indicating these neo-peptides were non-immunogenic. We took these data as the ground truth. Furthermore, the tumor-infiltrating TCRs of these 10 patients were obtained from RNA-seq data using MiXCR^44^, which is a universal tool for fast and accurate analysis of T- and B-cell receptor repertoire sequencing data. To this end, we used *TCRBagger* built in Section 4.2 to refine the rank of the candidate neopeptides based on the candidate neopeptide list identified by the three existing tools respectively, by further evaluating their immunogenicity by bagging the patient-specific TCR profiles. In this study, we also chose rank coverage scores (RCS) ^4^ to represent the performance of each tool to compare their patient-specific neoantigen prioritization performance, as applied in previous study ^4^ (**see Supplementary Methods**). The higher the RCS is, the better neoantigen prioritization performance.

As a result, compared to MuPeXI, the rank of known immunogenic neoantigens of 9 among 10 patients were improved, with the mean RCS increasing from -0.377 to 0.351 (**Fig.5e**). Compared to Neopepsee, the rank of known immunogenic neoantigens of 7 among 10 patients were improved, with the mean RCS increasing from 0.197 to 0.456 (**Fig.5f**). Compared to pTuneos, the rank of known immunogenic neoantigens of 6 among 10 patients were improved, with the mean RCS increasing from 0.346 to 0.411. Taking together, *TCRBagger* is demonstrated to be able to refine the rank list of candidate neopeptides and improve immunogenic neoantigen identification by interrogating the interaction between neopeptide and the patient-specific TCR profiles from tumor-infiltrating T cells under MIL (**Supplementary Data 5**).

## Discussion

The identification of neo-epitopes is critical in personalized tumor immunotherapy, providing a great opportunity to develop new vaccines, diagnostics and immunotherapies. As the peptide-TCR binding is the essential condition for neo-epitopes to trigger the T-cell response, an effective and robust neo-epitope identification model considering the peptide-TCR binding is in a high demand. Due to the inaccurate binding prediction for a single peptide-TCR pair and the lack of TCR binding information for most neo-epitopes, we developed *TCRBagger* to identify neo-epitopes by predicting their sample-specific TCR profiles using MIL from a bag-level perspective. *TCRBagger* is based on the simplified assumption that the binding of peptide-TCR is the essential condition for the neo-epitopes to trigger T-cell response, and the neo-epitopes are recognized or bound by at least one TCR in the T cell response while non-epitopes peptides are not recognized by any TCR.

There also exists limitations in *TCRBagger*, with further developments as follows. (1) In the current study, we only consider the beta chain of TCR in our model, as the TCR data with both alpha and beta chain information in existing databases is limited. However, *TCRBagger* is expected to be updated in the future when more information on alpha chains is available. (2) Currently *TCRBagger* did not consider the HLA typing information in the interaction modelling. The actual contributions of HLA typing in peptide-TCR binding recognition is important, while it remains an open question to be incorporated at present, considering that relatively few HLA typing information is available now. A previous study indicated that the effect of HLA type is small, as has been evaluated in peptide-TCR recognition in existing datasets. Furthermore, building models incorporating the exact HLA typing will limit the scope of the downstream analysis in current study. Nevertheless, the whole framework of *TCRBagger* can be easily extended by incorporating the encoding of HLA type in the future. (3) The current version of *TCRBagger* mainly focuses on the linear sequence of peptide-TCR interactions without considering the original three-dimensional structure of TCRs. Further studies can focus on the whole TCR three-dimensional structures when more structural data are available. (4) The current study is restricted on the HLA-I presented neo-epitope identification. Similar models can be built for HLA-II presented neo-epitope identification in the future. (5) *TCRBagger* would achieve a more robust performance with more available trainable data, consisting of sample-specific experimentally validated neo-epitopes and their infiltrating TCR profiles.

## Materials and Methods

### 1. Self-supervised pre-training for HLA-I restricted peptide representation

#### Using transformer encoder to obtain 512 features for each peptide

We collected a total of 16,055 HLA-I-restricted peptides after a strict filtering approach. To learn an efficient representation of HLA-I-restricted peptides, we applied the transformer encoder-based self-supervised learning and the mask language model (MLM) task (**Supplementary Fig.3**). We adopted a dynamic mask strategy with a 30% random mask rate for each amino acid in the training dataset and 15% in the validation dataset to evaluate the performance of our pre-trained model. Embedding dimensions, such as 64,128, 256, 512, were chosen in feature selection experiments aiming to select the best dimension (**Supplementary Fig.4a**). Four quantitative metrics, including training loss, validation loss, training masked amino acid accuracy and validation masked amino acid accuracy, were chosen to evaluate the performance of the self-supervised model with the same training epoch at 1,000. We found 512 embedding features achieving the best performance and then we chose as the peptide representation (**Supplementary Fig.4a**). It should be noted that our model tends to be stable at 30,000 training epochs with a validation loss of approximately 0.11 in the MLM task, which means that the global sequence information existing in each peptide was precisely extracted **(Supplementary Fig.4b**,**c**). Additionally, transformer encoder-based feature embedding instead of traditional one-hot encoding was adopted here because the former can obtain more distinct features when there is only one amino acid difference between two peptides, but their immunogenicity labels are different, which commonly existed in the current study. Of note, the aim of such representation study is to learn a proper peptide embedding used for the following CEWS analysis (**see Methods Section 2**), since the CEWS algorithm requires a highly accurate and efficient representation to characterize the similarity between two peptides.

The transformer encoder is made up of two main parts, including the multi-head attention layer and feedforward layer. Additionally, two residual connections are used in the transformer encoder, in which one is between the input layer and multi-head attention output layer and the other is between the output of the multi-head attention layer and the output of the feedforward layer. Our self-supervised pre-training model consists of three parts: (1) an amino acid embedding layer, (2) a sequence position encoding layer, and (3) six transformer encoder blocks. Compared to the traditional BERT model, the next sentence prediction (NSP) auxiliary task for pretraining, which has been indicated to have little effect on pretraining models, is avoided in our work. ^45–47^. We retain the mask language model (MLM) downstream auxiliary task. In each peptide sequence, 30% of random amino acids are masked in the training dataset, and 15% are masked in the validation dataset.

#### Self-attention mechanism in transformer block

The self-attention mechanism in the transformer encoder captures global amino acid information for each amino acid because of its ability to capture correlations between long-distance amino acids. Specifically, each sequence input can be represented as {*x*_1_, *x*_2_, …, *x*_*n*_}, where *x*_*i*_ denotes the *i*-th amino acid embedding. Therefore, the self-attention learns from the *Q, K, V* matrix and gives each amino acid pair, such as (*x*_1_, *x*_2_), an attention weight *α*, where the sum of *α* for each amino acid will be normalized to 1 by softmax.

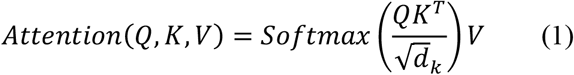

where *Q, K, V* here denote query, key, value matrix, respectively, 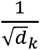 denotes the scaling factor.

Instead of using one-head self-attention, we used a multi-head self-attention mechanism that gives a peptide with different directions of attention. It has been proven that linearly projecting the *Q, K, V* matrices *h* times with various and learnable dimensions, *d*_*q*_, *d*_*k*_, *d*_*v*_ would be better without losing the cost of computation^29^.

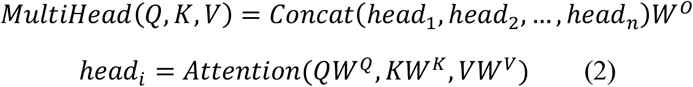

where 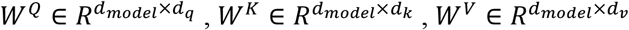 and 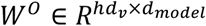 are the learnable parameters. *d*_*model*_ denotes the embedding dimension of each amino acid.

In this work, we set *d*_*model*_ to 512 and use 8 heads in the multi-head attention layer, so.*d*_*q*_ = *d*_*k*_ = *d*_*v*_ = *d*_*model*_/8.

#### MLM task

The mask language model (MLM) belongs to classical contextual self-supervised learning, aiming at extracting the context word information for a given word. Compared to traditional word embedding methods, such as word2vec, in which one word only has one embedding and less power to handle the polysemy problem^48^, the transformer-based language model begins with the word embedding matrix and position embedding matrix aiming to learn the dynamic word embedding at the last layer, which considers the ordering of words and the context of words^49^.

Given a peptide sequence *AAs* = {*a*_1_, *a*_2_, …, *a*_*n*_}, where *a*_*i*_ represents the *i*-th amino acid in this sequence, we can perform a similar process in NLP. We add the [*CLS*] token in front of the *AAs* and pad [*PAD*] tokens to each sequence until it achieves the fixed max length, which is 16 in this work. In the training process, each amino acid has a 30% chance of being replaced by a [*MASK*] token in the training dataset and a 15% chance in the validation dataset. To mitigate the mismatch problem and improve the performance, we replace these amino acids with [*MASK*] 80% of the time, replace these amino acids with random amino acids 10% of the time and keep these amino acids unchanged 10% of the time^49^. Then, the MLM task is designed to predict the ground truth of the [*MASK*] token. Therefore, our mission is to minimize the loss function as follows:

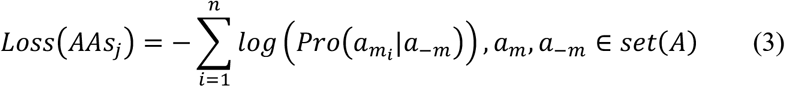

where *AAs*_*j*_ represents the input peptide *j* sequence. 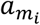 is the *i*-th masked amino acid in peptide *AAs*_*j*_, and *a*_−*m*_ denotes the amino acids not masked amino acids in peptide *AAs*_*j*_. *A* is the set of amino acids.

In the model training process, we set the maximum learning rate as 0.001, the minimum learning rate as 1 × 10^−5^, and the warmup proportion as 0.1, and we trained the model at 30,000 epochs.

### 2. Denoising learning for inaccurately experimentally annotated peptides based on self-supervised learned peptide embedding

In the setting of inaccurately supervised learning, identifying and handling mislabeled instances is important. The knowledge learned from the model is very sensitive to the perturbation of noisy data, which means that mislabeled instances or noisy data may heavily affect the model’s generalization. Therefore, the cut edge weight statistic (CEWS) algorithm was proposed to handle this problem, which has been proven to enhance the classification accuracy^50^. Based on the discovery that the immunogenicity state of peptides was inconsistent in the two databases, we rectified the peptide labels used in our model. The essence of this algorithm can be generalized as the following three main ideas: First, construct the weighted neighborhood graph matrix. Then, identify the cut edge between data points. Finally, the sum of cut edge weights is calculated, and a two-sided hypothesis test is applied to identify mislabeled instances. Of note, self-supervised learned peptide embedding is used in this step. An additional ablation study is presented to indicate the necessity to use the learned peptide embedding rather than the Atchely factor based peptide embedding for denoising study **(See supplementary methods, Supplementary Fig.5)**.

#### Weight matrix calculation

To measure the degree of proximity between peptides, we use Euclidean distance to calculate the distance between peptide *i* and peptide *j*. Then, if there is an edge satisfying the following conditions, peptide *i* and peptide *j* will have an edge. The edge weight between them can be formulized as well.

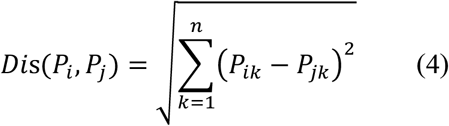

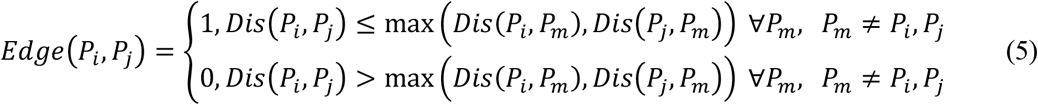

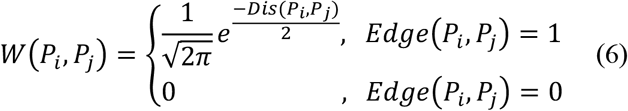

where *P*_*i*_ is peptide *i* and *n* is the length of the peptide representation vector. *Edge*(*P*_*i*_, *P*_*j*_) denotes whether there is an edge between peptide *i* and peptide *j*. 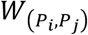 denotes the weight between peptide *i* and peptide *j* if they have an edge between them.

#### Cut edge definition

If peptide *i* and peptide *j* are connected while they have different labels, then the edge between them will be defined as the cut edge. Intuitively, for each peptide, if it has more cut edges in its surrounding neighbors, it will have a higher probability of being a mislabeled instance.

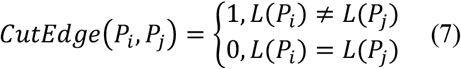

where *CutEdge*(*P*_*i*_, *P*_*j*_) denotes whether peptide *i* and peptide *j* possess different associated labels and *L*(*P*_*i*_) denotes the immunogenicity state of peptide *i*.

#### J-value statistic calculation

The prior probability of a peptide labeled positive or negative is calculated separately, which is commonly estimated by the fraction of positive (negative) instances in the data domain. After that, the label confidence estimation of each peptide can be calculated with *J* − *value*.

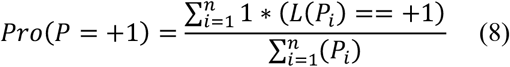

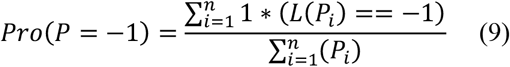

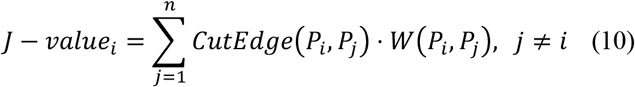

where *Pro*(*P* = *+*1) denotes the empirical probability of an instance labeled positive and *Pro*(*P* = −1) denotes the empirical probability of an instance labeled negative. *J* − *value*_*i*_ denotes the label confidence of a peptide *i*.

Therefore, we hypothesize *H*_0_ that the instances in the neighborhood graph we constructed above are labeled independently with probability *Pro*(*P* = *+*1) and *Pro*(*P* = − 1). Then, each edge with respect to the peptide *i* should correspond to a Bernoulli distribution with 1 indicating a cut edge and 0 otherwise. Thus, the *Pro*(*CutEdge*(*P*_*i*_, *P*_*j*_) = 1) should be equal to the 1 − *Pro*(*P*_*i*_ = *L*(*P*_*i*_)). When the data size is large enough, *J* − *value* can be modeled by the normal distribution. Therefore, we can use *μ*0 and *σ*^2^ to normalize them to the standard normal distribution *N*(0,*m*). The hypothesis test can be performed to ℂ. Based on the assumption above, we can conclude that the smaller ℂ has higher confidence about its label, and vice versa. We selected the strict *α* = 0.01 as the rejection threshold in our project, and the *RejectionDomain* of the peptides can be calculated.

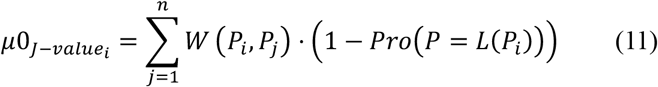

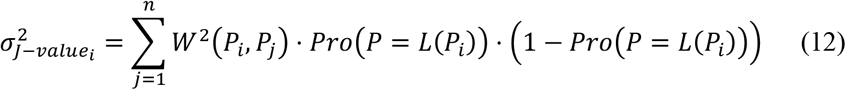

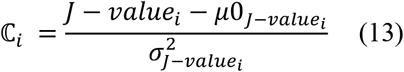

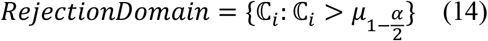

where 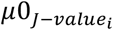 denotes the expected *J* − *value* of peptide *i* and 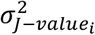 denotes the variance in the *J* − *value* of peptide *i*. ℂ_*i*_ denotes the *J* − *value* of peptide *i* normalized into the standard normal distribution, which can represent whether peptide *i* is mislabeled. *RejectionDomain* denotes the set of mislabeled peptides.

### 3. Multi-instance learning used in *TCRBagger*

#### Model assumption

The proposed model is inspired by a kind of weakly supervised learning, i.*e*., a multi-instance learning model^19^. It shows the concept that a bag containing as long as one key component will be a positive bag^51^. In our study, since the binding of pMHC - T cell receptor TCR is the essential condition for neo-epitope to trigger immune response, *TCRBagger* is designed to identify neo-epitope by predicting their binding TCR profiles from a bag-level perspective, based on the simplified assumption that the immunogenic peptides(neo-epitopes) are recognized or bound by at least one TCR in the T cell response while non-immunogenic peptides are not recognized by any TCR. Specifically, peptide is represented by bagging of a corresponding sample-specific TCR repertoire. A bag *X* is made up of a peptide *P* and TCR repertoire having plenty of TCRs {*TCR*_1_, *TCR*_2_, …, *TCR*_*n*_}. Then, a bag can be represented by *X*_*k*_ = {*x*_1_, *x*_2_, …, *x*_*n*_}, where *x*_*i*_ indicates a *P*_*k*_ − *TCR*_*i*_ pair. Furthermore, we assume the hidden binding labels of *x*_*i*_ is *y*_*i*_, indicating the peptide and TCR are binding or not, which we do not know in the training process. If there is at least one binding TCR within one bag *X*_*k*_, the immunogenicity state *Y*_*k*_ of bag *X*_*k*_ will be positive. Otherwise, the immune state *Y*_*k*_ of bag *X*_*k*_ will be negative. Therefore, the multi-instance learning problem can be formulized as follows:

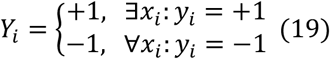

MIL classification problems are mainly divided into two domains: bag level and instance level^19^. In our study, we choose bag-level embedding to classify the immunogenicity state *Y*_*k*_ of peptide *P*_*k*_, as it has a better performance in error tolerance, for which some mislabeled negative instances within positive bags may not influence the final prediction result. Furthermore, we adopt bag accuracy as an optimization criterion because we mainly focus on the peptide immunogenicity state where the general performance of a peptide to a TCR repertoire is most significant. The assumption above also implies that the instances within a bag must be permutation-invariant, which means there is no ordering between them.

#### The model architecture of *TCRBagger*

The property of a bag that is filled with multiple instances and each instance presented by a fixed-length vector allows it to be featured as an *M* × *N* matrix, where *M* indicates the number of bags and *N* indicates the number of instances. Then we use a convolutional layer as the base feature extractor. The CNN architecture has been widely used in the field of image recognition and has made considerable progress. Therefore, the main architecture of the *TCRBagger* is based on the CNN and gate attention mechanism. The gate attention mechanism guarantees a learnable pooling layer that can improve the integration performance of instances and be able to have a relatively better interpretation for our model. As the instance number within one bag can be different, we set up 100 as the max limitation of instance number. Therefore, bags with instance numbers less than 100 will be handled with padding and masking operations when they are input into our model. Meanwhile, instance numbers in bags larger than 100 will be divided into multiple bags, where the bag with the highest immunogenicity score will be chosen as the eventual bag score. In our model architecture, we defined 32 convolution kernels with 1 × 4 dimensions in the first and second convolutional layers. Following the max-pooling layer with a 1 × 2 kernel, dropout layers with a drop rate of 0.5 were stacked. The activation function *Relu* was applied to each convolutional layer. After extracting the feature of each instance based on the conventional layers, each bag can be transformed into 100 × *N*_*l*_ × 32. To meet the requirement of the permutation-invariant property, we reshape the output matrix to 100 × *N*_*new*_, where *N*_*new*_ = *N*_*l*_ × 32, so that each instance can be represented by *m* × *N*_*new*_. Then, a fully connected layer with 128 neural nodes was applied to the output matrix, and the representation of each instance translated into 128 dimensions, which we called instance deep representation. (**Supplementary Fig.6**). Furthermore, the gate attention mechanism-based attention block was applied to each bag for general bag embedding. After that, the final bad embedding was taken to calculate the loss with ground truth.

#### Attention mechanism in *TCRBagger*

Compared with traditional pooling methods such as max pooling and mean pooling, we used the weighted average instances while each weight of instance was obtained while training. Meanwhile, the sum of each weight is normalized to be 1 by softmax activation. Then, the attention base MIL pooling is listed below:

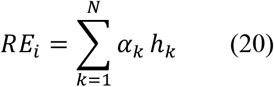

where

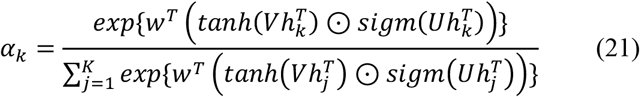

*α* is set based on the gated attention mechanism, which gives a learnable nonlinearity, avoiding the troublesome linearity in *tanh*. where *w* ∈ *R*^*L*×1^ and *V, U* ∈ *R*^*L*×*M*^ are trainable parameters. *RE*_*i*_ refers to the bag embedding of *X*_*i*_, and *h*_*k*_ refers to the deep representation vector of instance *k*.

#### Instance embedding of *TCRBagger*

To construct a bag, we need to model each instance in the bag so that all instance embeddings are integrated as bag embedding. Concretely, we need a fast and efficient way to depict the binding features of peptides and TCRs. First, each amino acid with the chemical properties of hydrophilia, hydrophobicity and so on was extracted and represented as 5-dimensional features, *A*_*i*_ ∈ *R*^5×1^, by Atchley factors. To precisely calculate the binding features of the peptide and TCR, we used 3 to 5 fragments of TCR to slide it along the peptide sequence, which has been proven to have a higher affinity, and amino acid pairwise contact potential (AACP) scales were used. Then, we calculated the statistical features of the results from the sliding TCR fragment as the binding features of peptide-TCR pairs. This calculation method is the basic idea of the ‘global-local’ algorithm, which is depicted below. Therefore, the instance embedding feature can be expressed as the combination of binding features and amino acid property features. Of note, in the MIL framework, we avoid to use the former self-supervised learned embedding for peptide representation, although it is highly accurate in the representation of peptide itself, it is not efficient in the modeling of the interaction between peptide and TCR profiles.

Given a peptide *P*_*i*_ and a TCR sequence *T*_*i*_ in which the head and tail 3 amino acids are dropped, we *Split* the peptide and TCR sequence with respect to the fragment length into multiple fragments. We then calculated the binding features *B* of each pair of fragments between the peptide and TCR. For each TCR fragment, we will choose the maximum of its binding values to the peptide sequence as its representation. Then, we fully use some statistics to capture the information from all representations *S*_*k*_ for each fragment length, *F*_*k*_.

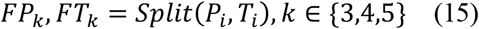

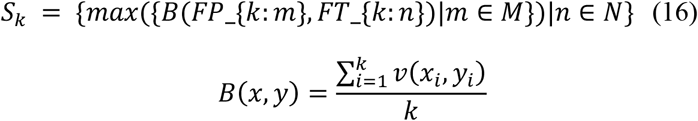

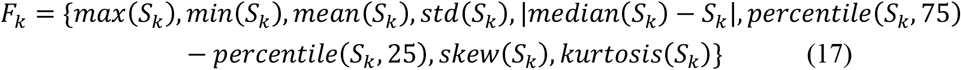

where *FP*_*k*_, *FT*_*k*_ denotes the collection of peptide fragments and TCR fragments for a specific fragment length.

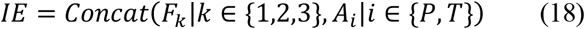

where *IE* denotes an instance embedding that combines the binding features and amino acid property features.

#### Training loss of *TCRBagger*

Given a set of training peptides and their corresponding TCR repertoires *𝒯* = {*X*_1_, *X*_2_, …, *X*_*n*_}, with their ground truth immune state *𝒴* = {*Y*_1_, *Y*_2_, …, *Y*_*n*_}, our model aims to minimize the loss function:

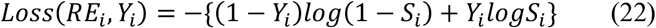

where *RE*_*i*_ is the gate attention-based bag embedding score and *Y*_*i*_ is the bag label corresponding to the TCR repertoire.

All parameters of our multi-instance model are trainable, and we use the *Adam* optimizer in TensorFlow with a learning rate of 0.001 to train out the model by backpropagation.

#### Attention visualization of *TCRBagger*

To illustrate the interpretation of our model, we visualize the attention block in our model in which the gate attention weight for each TCR will be calculated. Therefore, we use our model to predict the bags in our validation set and extract the attention block in them. The attention block was visualized by the *m*0 × *m*0 cell matrix, where each cell represents a TCR attention weight. The numbers 1 and 0 in the cell denote whether the TCR will have a higher response probability with the peptide in this bag. As the sum of attention weights {*α*_1_, *α*_2_, …, *α*_100_} was normalized to 1 by softmax, the color shade represents the degree of attention magnitude in the cells, where a deeper color means a higher attention weight, and vice versa.

## Supplementary Information

**Supplementary Methods 1-4, Supplementary Table 1 and Supplementary Figs.1-6**

## Supplementary Data

Supplementary Data 1: The filtered neo-epitopes and their binding TCRs.

Supplementary Data 2: The neo-epitope confidence score of each filtered peptide, which estimated by the CEWS algorithm.

Supplementary Data 3: Seen dataset and Unseen dataset used for evaluate three single TCR-peptide interaction prediction methods.

Supplementary Data 4: The integrated table of peptides and TCRs from six independent single-cell level experiments.

Supplementary Data 5: The summary table of neoantigens predicted by the three softwares and their rectified priority given by *TCRBagger* in project PRJNA243084, PRJNA278450, PRJNA298310, PRJNA298330.

## Acknowledgements

Not applicable

## Authors’ contribution

Liu Qi designed the framework of this work. Yuli Gao and Yicheng Gao performed the pipeline design and analyses. Wannian Li, Qisi Wu, Feiyang Xing, Chi Zhou, Shaliu Fu, Guohui Chuai, He Zhang performed the data curation and data analysis. Yuli Gao, Yicheng Gao and Liu Qi wrote the manuscript with the help of other authors. All authors read and approved the final manuscript.

